# Novel *trans*-translation-associated gene regulation revealed by prophage excision-triggered switching of ribosome rescue pathway

**DOI:** 10.1101/2022.04.27.489667

**Authors:** Haruka Onodera, Tatsuya Niwa, Hideki Taguchi, Yuhei Chadani

**Affiliations:** School of Life Science and Technology, Tokyo Institute of Technology, S2-19, Nagatsuta 4259, Midori-ku, Yokohama, 226-8503, Japan; Cell Biology Center, Institute of Innovative Research, Tokyo Institute of Technology, S2-19, Nagatsuta 4259, Midori-ku, Yokohama, 226-8503, Japan

**Keywords:** *Escherichia coli*, ribosome rescue, *trans*-translation, phage regulatory switch, SsrA RNA, ArfA, polynucleotide phosphorylase (PNPase), prophage, upstream ORF, downstream ORF

## Abstract

*Escherichia coli* has multiple pathways to release a nonproductive ribosome complex stalled at the 3’ end of nonstop mRNAs: SsrA RNA-mediated *trans*-translation and stop codon-independent termination by ArfA/RF2 or ArfB (YaeJ). The *arfA* mRNA lacks a stop codon and thus its expression is repressed by *trans*-translation. Therefore, ArfA is considered to complement the ribosome rescue activity of *trans*-translation, but the situations in which ArfA is expressed to rescue the nonproductive complexes have not been elucidated. Here, we demonstrated that the excision of the CP4-57 prophage adjacent to the *E. coli ssrA* gene leads to the inactivation of SsrA RNA and switches the primary rescue pathway from *trans*-translation to the ArfA/RF2 pathway. A comparative quantitative proteomic analysis revealed that the switching of the rescue pathway rearranges not only the proteome landscape in *E. coli* cells but also the phenotype, such as motility. Among the proteins with significantly increased abundance in the *ssrA*-inactivated cells, we found ZntR, whose mRNA is transcribed together as the upstream part of the nonstop *arfA* mRNA. Further analysis revealed that the translation of the nonstop ORF of *arfA* triggered the <u>**r**</u>epression of the <u>**u**</u>pstream *zntR* ORF, via ***<u>t</u>****rans*-translation-coupled <u>**ex**</u>onucleolytic (RUTEX) degradation by a polynucleotide phosphorylase. These results provide a novel example of *trans*-translation-dependent regulation, and shed new light on the physiological roles of prophages in gene expression.

## Introduction

Ribosomes translate mRNA from the start codon to the stop codon for protein synthesis. The start codon and the surrounding nucleotide sequences, such as the Shine-Dalgarno sequence, define the entry site of the 30S initiation complex (1). The stop codon is indispensable for the hydrolysis of the peptidyl-tRNA within the ribosome by release factors (RF) and the following subunit dissociation from the mRNA (1). However, the translation of an aberrant mRNA lacking an in-frame stop codon traps the ribosome at its 3’ end, due to the defect in the termination reaction. Such stop codon-less mRNAs (nonstop mRNA) are constantly generated by mRNA degradation and premature transcription termination, and waste at least 2-4 % of total translation (2, 3). Therefore, recycling of the ribosomes stalled on the aberrant mRNA is essential to maintain the translation activity and cell viability. In fact, multiple ribosome rescue pathways are harnessed to resolve this problem in various bacterial species.

Most stalled ribosomes are rescued by SsrA RNA (tmRNA) and SmpB-mediated *trans*-translation (4, 5). The *ssrA* gene is highly conserved among all bacterial genomes determined to date (6, 7), implying the physiological importance of *trans*-translation. SsrA RNA has both the tRNA-like domain, which can accept alanine, and the mRNA-like domain containing an in-frame stop codon. After the SsrA RNA enters the vacant A-site of the stalled ribosome, the ribosome switches the reading frame from the nonstop mRNA to the mRNA-domain of SsrA RNA to properly terminate the translation. This results in a truncated polypeptide translated from the nonstop mRNA (hereafter called a “nonstop polypeptide”) fused with the SsrA RNA-encoded degradation tag at its C-terminus, which promotes degradation by proteases such as ClpXP (8). *Trans*-translation also induces the rapid degradation of nonstop mRNA (9). Previous research using artificial constructs showed that RNase R, one of the major exonucleases in *E. coli*, degrades the nonstop mRNA in a *trans*-translation-dependent manner (10). Thus, *trans*-translation is a sophisticated mechanism that rescues the stalled ribosome and consequently avoids the accumulation of aberrant proteins and mRNAs.

Despite several lines of evidence supporting the importance of *trans*-translation, many bacteria, such as *Escherichia coli, Francisella tularensis, Bacillus subtilis* and *Caulobacter crescentus*, can survive even in the absence of the SsrA RNA (11–14). Some of these bacteria reportedly possess alternative ribosome rescue pathways (15–17). ArfA (alternative ribosome rescue factor A) in *E. coli* cooperates with RF2 to terminate translation by hydrolyzing the peptidyl-tRNA in a stop codon-independent manner (18). In contrast to *trans*-translation, the ArfA/RF2 pathway does not add a degradation tag to the released polypeptide and apparently does not recruit exonucleases to the nonstop mRNA. Therefore, nonstop polypeptides could accumulate if the stalled ribosome is released by ArfA (19).

ArfA expression is tightly repressed by multiple regulations (20, 21). A stem-loop structure, which functions as both a rho-independent terminator and a recognition site for RNase III, is encoded within the *arfA* ORF (20, 21). Therefore, the *arfA* mRNA is expressed as a nonstop mRNA in various situations and is repressed by the SsrA RNA-mediated *trans*-translation. Moreover, even if the ribosome fortuitously terminates the translation of the *arfA* ORF at the endogenous stop codon, the degradation of the full-length ArfA protein is induced by the hydrophobic amino acid cluster at the C terminus (21). Therefore, ArfA is strongly repressed under the conditions where *trans*-translation is active. In other words, ArfA likely functions as a backup mechanism in situations where the *trans*-translation activity is somehow disturbed. Alternative ribosome rescue factors in other bacterial species are also repressed by the SsrA RNA, indicating that this regulation is commonly beneficial to bacterial species (17). However, the situations in which the alternative ribosome rescue pathway is activated are poorly understood.

A recent study showed that the excision of the CP4So prophage, adjacent to the *ssrA* gene in the *Shewanella oneidensis* genome, induces a single nucleotide deletion in the *ssrA* gene and abolishes its function (22). Such alteration of the expression/function of the host genes by prophage excision has been referred to as a “phage regulatory switch” in previous reports (23–25). The *E. coli* K-12 strain also has a prophage (CP4-57) downstream from the *ssrA* gene (26). Although CP4-57 excision rarely occurs under normal conditions, the excision rate increases to 1-2 per 10,000 cells in specific situations, such as biofilms or long-term cultivation (27, 28). The excision of CP4-57 reportedly causes the deletion of T357 in *ssrA*, which forms a wobble base pair in the acceptor stem in the tRNA-like domain, as in *S. oneidensis* (28) (**Fig. S1A**). However, the influences of this mutation on the SsrA RNA activity and the activation of an alternative ribosome rescue pathway have not been well documented.

Based on these previous findings, we examined the possibility that CP4-57 excision switches the primary ribosome rescue pathway from SsrA RNA-mediated *trans*-translation to ArfA/RF2-mediated termination, by impairing the SsrA RNA activity. In this report, we found that (i) CP4-57 excision impairs the *trans*-translation and switches the ribosome rescue pathway from *trans*-translation to the ArfA/RF2 pathway, and (ii) CP4-57 excision rearranges the cellular proteome and the phenotype in a *trans*-translation-dependent manner. Further analysis revealed that *zntR*, which is transcribed as a polycistronic mRNA with *arfA*, is repressed in a *trans*-translation-dependent manner. The *zntR* repression is independent of proteolysis, but dependent on *trans*-translation-coupled exonucleolytic degradation by a polynucleotide phosphorylase (PNPase), indicating that the degradation of the *zntR* region in the mRNA is associated with that of the *arfA* nonstop mRNA. These results revealed a novel mode of *trans*-translation-dependent regulation, in that the expression of the upstream ORF followed by the nonstop ORF is also influenced by the ribosome rescue pathways.

## Results

### The excision of CP4-57 prophage switches the ribosome rescue pathways

Based on several previous reports, we hypothesized that the CP4-57 excision-triggered *ssrA*ΔT357 mutation inactivates *trans*-translation and consequently switches the ribosome rescue pathway to the alternative ArfA/RF2 pathway (**Fig. 1A**). To evaluate the *trans*-translation activity of the SsrAΔU357 mutant, we examined whether SsrA-tagged polypeptides are accumulated in cells. Since SsrA-tagged polypeptides are rapidly degraded, we used SsrA^His^, an SsrA RNA variant encoding a proteolysis-resistant hexahistidine tag (29) (**Fig. 1B**). Cellular extracts of *ssrA* deletion mutants expressing SsrA^His^ or SsrA^His^ΔT357 were analyzed by western blotting, using an anti-His_6_ antibody **(Fig. 1C**). In the presence of SsrA^His^, numerous polypeptides with various molecular weights were detected, indicating the *trans*-translation reaction by the SsrA^His^ mutant, as reported previously (29). In contrast, the expression of SsrA^His^ΔU357 did not produce any polypeptide containing the His_6_-tag, indicating that the ΔU357 mutation disrupts the *trans*-translation activity of the *E. coli* SsrA RNA. Note that the possibility of SsrA inactivation by prophage excision has been suggested for other γ-proteobacteria that possess a homolog of *arfA* (22, 28, 30–32) (**Table 1**). Accordingly, we introduced the mutations found in those bacteria into the *E. coli* SsrA RNA, and confirmed the robust inactivation of the *trans*-translation activity, which was similar to the ΔU357 mutation (**Fig. S1B lanes 6 and 7**).

**Fig. 1.**
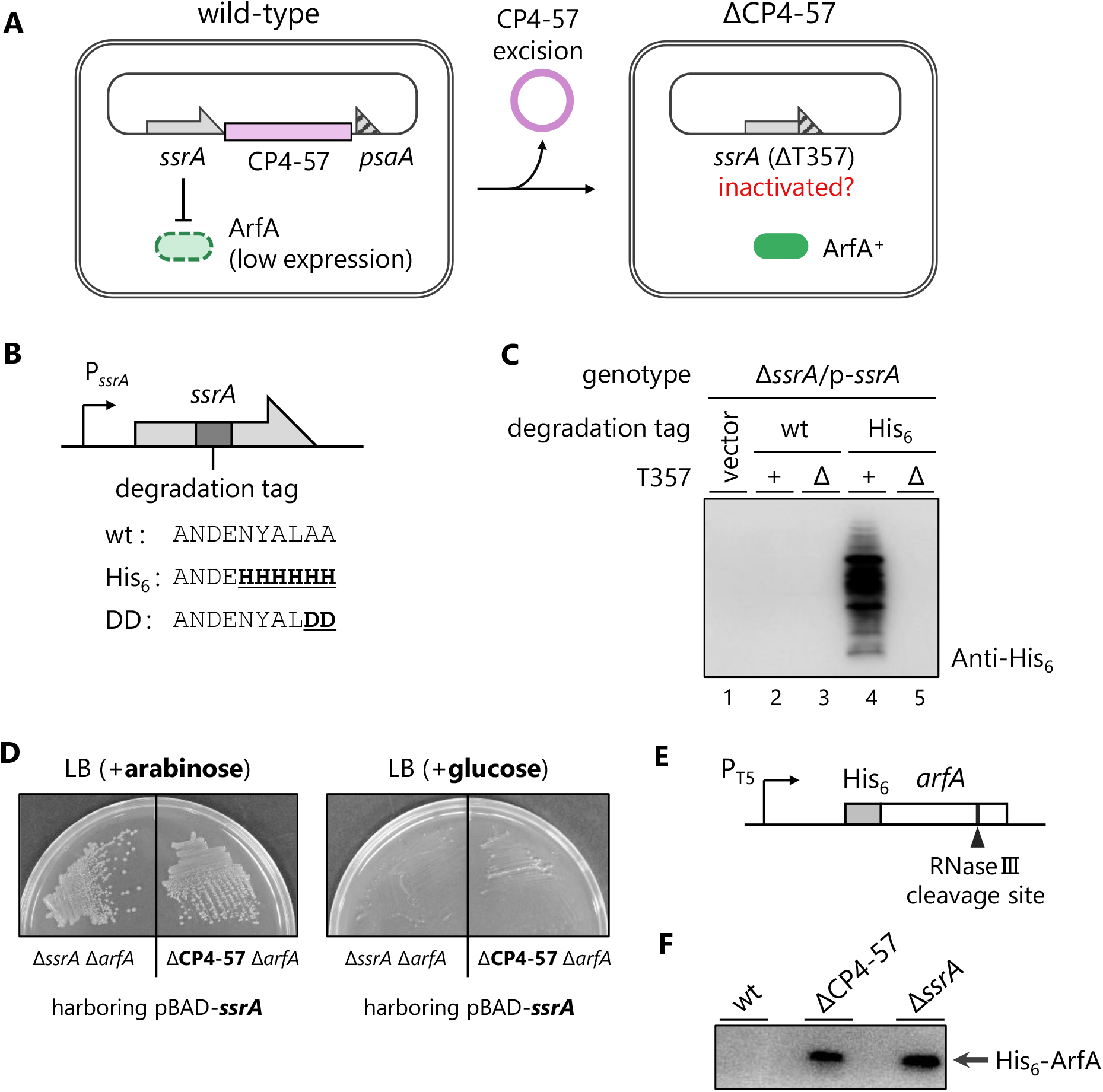
CP4-57 prophage excision is a junction point to switch the primary ribosome rescue pathway in *E. coli*. (A) Working hypothesis: the CP4-57 excision-induced *ssrA*ΔT357 mutation irreversibly inactivates *trans*-translation and subsequently activates the ArfA/RF2 alternative rescue pathway. (B) Schematic of the plasmids expressing SsrA RNA, and amino acid sequences of the degradation tags encoded within the wild-type SsrA RNA and its variants, SsrA^His^ and SsrA^DD^. (C) Visualization of aberrant polypeptides rescued by *trans*-translation. The *E. coli* Δ*ssrA* strain harboring pMW118 (vector control, lane 1), p-*ssrA* (lane 2), p-*ssrA*ΔT357 (lane 3), p-*ssrA*^His^ (lane 4) or p-*ssrA*^His^ΔT357 (lane 5) was grown in LB medium until mid-log growth phase. The cells were then collected, extracted, fractionated by SDS-PAGE, and analyzed by western blotting using an anti-His_6_ antibody. (D) Synthetic lethal phenotype of ΔCP4-57 and Δ*arfA* mutation. *E. coli* Δ*ssrA* Δ*arfA* and the ΔCP4-57 Δ*arfA* strain harboring pBAD-*ssrA* were grown on LB plates containing 0.2 % arabinose (left) or 0.4 % glucose (right) at 37 °C overnight. (E) Schematic of the plasmid (pCH200 (15)) expressing N-terminally His_6_-tagged ArfA. The *arfA* ORF and the His_6_-tag sequence are shown by open and filled boxes, respectively. The IPTG-inducible LacO-T5 promoter (P_T5_) and the RNase III cleavage site are also shown. (F) Expression profile of His_6_-ArfA in wild-type, ΔCP4-57 and Δ*ssrA* strains. Each strain harboring pCH200 was cultured in LB medium until the OD_660_ reached 0.2-0.3. The expression of ArfA was then induced for 2 hrs in the presence of 500 µM IPTG. The extracts of each strain were analyzed by western blotting using an anti-His_6_ antibody.

**Table 1.**
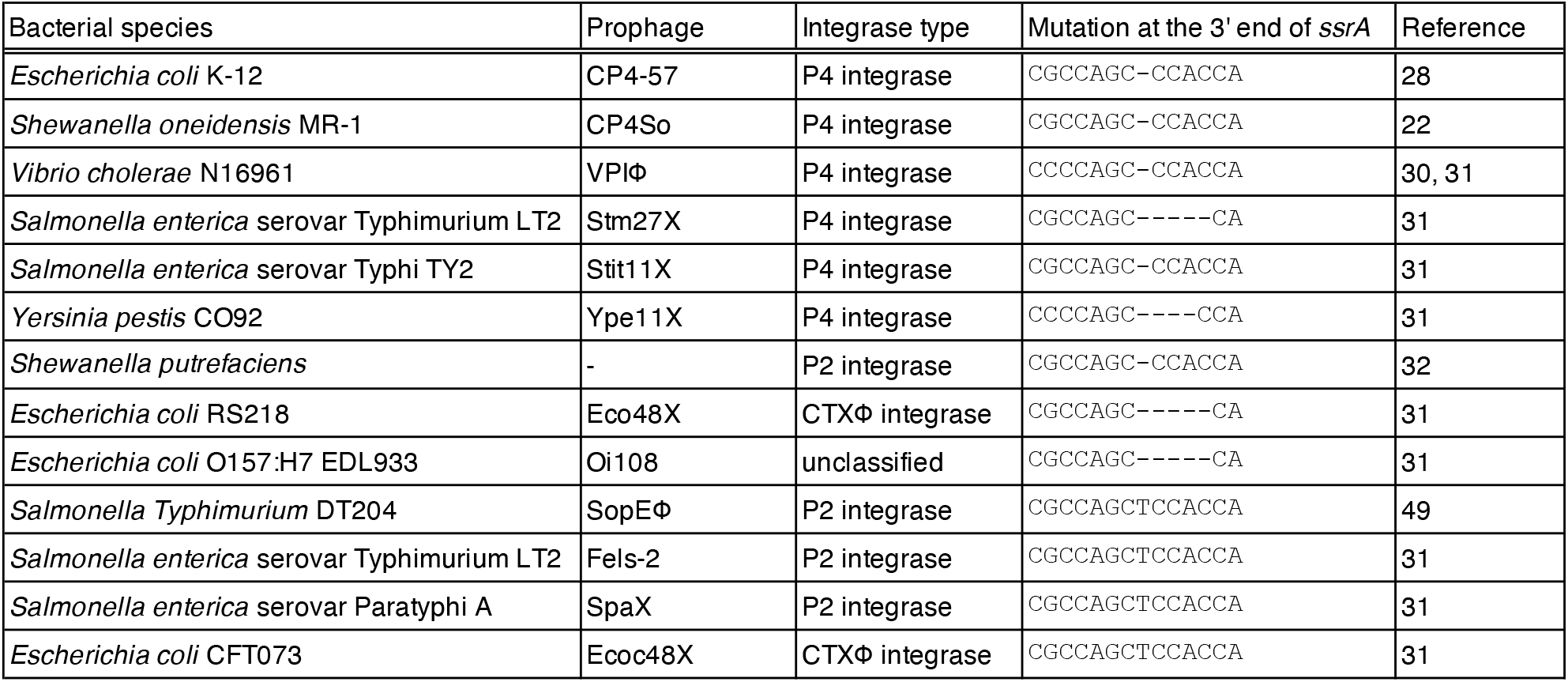
Prophage excision-induced mutations in *ssrA* gene among bacterial species.

| Bacterial species | Prophage | Integrase type | Mutation at the 3' end of *ssrA* | Reference |
| --- | --- | --- | --- | --- |
| *Escherichia coli* K-12 | CP4-57 | P4 integrase | CGCCAGC-CCACCA | 28 |
| *Shewanella oneidensis* MR-1 | CP4So | P4 integrase | CGCCAGC-CCACCA | 22 |
| *Vibrio cholerae* N16961 | VPIΦ | P4 integrase | CCCCAGC-CCACCA | 30, 31 |
| *Salmonella enterica* serovar Typhimurium LT2 | Stm27X | P4 integrase | CGCCAGC-----CA | 31 |
| *Salmonella enterica* serovar Typhi TY2 | Stit11X | P4 integrase | CGCCAGC-CCACCA | 31 |
| *Yersinia pestis* CO92 | Ype11X | P4 integrase | CCCCAGC----CCA | 31 |
| *Shewanella putrefaciens* | - | P2 integrase | CGCCAGC-CCACCA | 32 |
| *Escherichia coli* RS218 | Eco48X | CTXΦ integrase | CGCCAGC-----CA | 31 |
| *Escherichia coli* O157:H7 EDL933 | Oi108 | unclassified | CGCCAGC-----CA | 31 |
| *Salmonella Typhimurium* DT204 | SopEΦ | P2 integrase | CGCCAGCTCCACCA | 49 |
| *Salmonella enterica* serovar Typhimurium LT2 | Fels-2 | P2 integrase | CGCCAGCTCCACCA | 31 |
| *Salmonella enterica* serovar Paratyphi A | SpaX | P2 integrase | CGCCAGCTCCACCA | 31 |
| *Escherichia coli* CFT073 | Ecoc48X | CTXΦ integrase | CGCCAGCTCCACCA | 31 |

We previously reported that *E. coli* cells lacking both SsrA RNA and ArfA are unable to grow due to the accumulation of nonstop translation complexes (15). Therefore, we examined whether the deletion of *arfA* and the CP4-57 excision (ΔCP4-57 *ssrA*ΔT357) also elicit a synthetic lethal phenotype. The CP4-57 excision was induced by the overexpression of the AlpA protein, as described previously (27, 28). The *arfA* gene was then deleted in the presence of pBAD-*ssrA*, which produces the SsrA RNA in response to arabinose (15). As expected, the ΔCP4-57 Δ*arfA* strain, as well as the Δ*ssrA* Δ*arfA* double mutant, grew only in the presence of arabinose (15) (**Fig. 1D**). This result further confirmed that SsrAΔU357 impairs the *trans*-translation activity.

Finally, we investigated whether the ribosome rescue pathway is actually switched from *trans*-translation to the ArfA/RF2 pathway in the ΔCP4-57 strain. We expressed *arfA* encoding the N-terminal His_6_-tag and the stem-loop for RNase III cleavage in *E. coli* strains (**Fig. 1E and 1F**). The His_6_-ArfA protein did not accumulate in the wild-type strain, due to the tight repression by *trans*-translation (20, 21). In contrast, the ArfA protein was expressed in the ΔCP4-57 strain and the Δ*ssrA* mutant. This result clearly demonstrated that the ArfA/RF2 pathway is the primary system to rescue the nonstop translation complex in the ΔCP4-57 strain. From these results, we concluded that the excision of *E. coli* CP4-57 functions as a phage regulatory switch to convert the primary ribosome rescue pathway from SsrA RNA-mediated *trans*-translation to the ArfA/RF2 system.

### CP4-57 excision rearranges the *E. coli* proteome by switching the ribosome rescue pathways

Both SsrA RNA and ArfA/RF2 rescue stalled ribosomes; however, only the former pathway induces the degradation of both the released polypeptide and the nonstop mRNA (15, 21). Therefore, we assumed that the CP4-57 excision apparently shuts off the clearance mechanism for nonstop mRNAs and their translational products. This would result in the accumulation of nonstop polypeptides and thus rearrange the proteome landscape. To validate this possibility, we performed a quantitative proteomics analysis by using the SWATH-MS acquisition method (33). We compared the *E. coli* BW25113 standard strain (wild-type) and its derivatives, ΔCP4-57, Δ*ssrA*, Δ*intA*-*ypjF* (lacking the genes within the CP4-57 prophage region while retaining *ssrA*) and Δ*ssrA*-*ypjF*, as summarized in **Fig. 2A**. Each strain was cultivated in LB broth until mid-log phase and then processed for the comparative analyses. The fold change values of each protein were calculated as the division of the MS-intensity in the mutant by that in the wild-type standard strain. Volcano plots revealed the dynamic rearrangement of the proteome landscape in the ΔCP4-57, Δ*ssrA* and Δ*ssrA*-*ypjF* mutants (**Fig. 2B**). Interestingly, the regulons of the heat shock transcriptional regulator, σ32, were upregulated in the SsrA-deficient cells (34) (**Fig. S2A**). This result is consistent with the previous report that the heat shock response is caused by the accumulation of aberrant polypeptides in the Δ*ssrA* mutant (35). In contrast to the other mutations, the Δ*intA*-*ypjF* deletion poorly affected the proteome landscape under our test conditions.

**Fig. 2.**
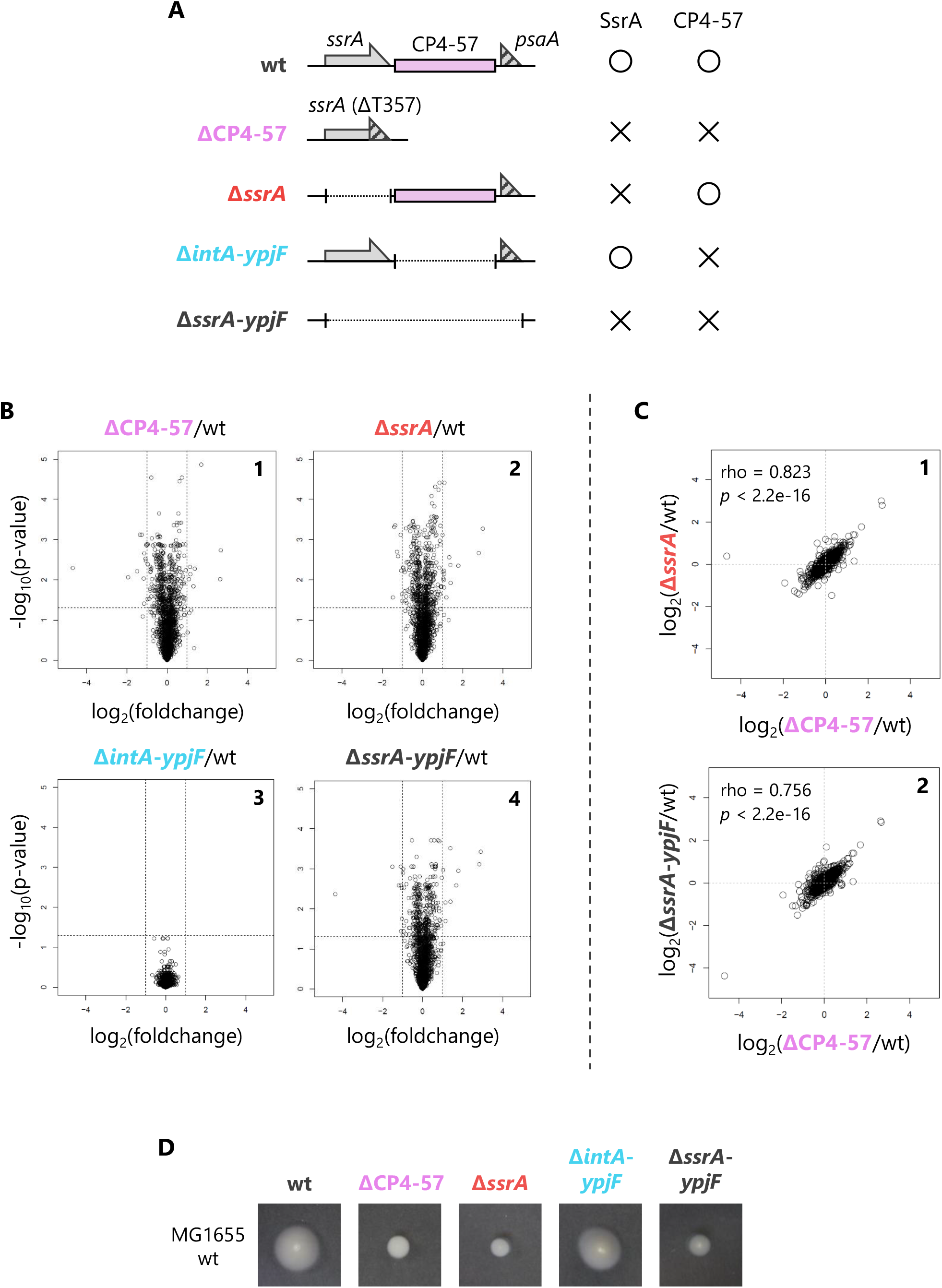
CP4-57 excision rearranges the proteome landscape in *E. coli* cells by switching the primary ribosome rescue mechanism. (A) Schematic drawing of the genomic structure of the *ssrA* - CP4-57 locus in the *E. coli* BW25113 strain (wild type: wt) and its derivatives, ΔCP4-57, Δ*ssrA*, Δ*intA*-*ypjF* (lacking the entire sequence of CP4-57 prophage region while retaining intact *ssrA*) and Δ*ssrA*-*ypjF. E. coli* cells were grown in LB medium until mid-log phase (OD_660_ ∼0.5) and collected for the SWATH analysis (See “Materials and Methods” for details). (B) The degrees of proteomic rearrangement between BW25113 (wt) and its derivatives (ΔCP4-57: panel 1, Δ*ssrA*: panel 2, Δ*intA-ypjF*: panel 3, Δ*ssrA*-*ypjF*: panel 4) are represented by volcano plots. Each dot represents the fold change and the *P*-value of each protein identified by the SWATH-MS analysis, plotted according to its relative abundance ratio (log_2_ fold change) on the horizontal axis and *P*-value on the vertical axis. The lines indicate a *P*-value of 0.05 and 2-fold change. (C) Two-dimensional plots of the fold change values in the *E. coli* BW25113 mutant are indicated below and on the side of the graph. Each plot is represented with Spearman’s rho and *P*-values calculated by Spearman’s rank correlation tests. (D) Swimming motility of MG1655 wild-type strain and its derivatives, the ΔCP4-57, Δ*ssrA*, Δ*intA*-*ypjF* and Δ*ssrA*-*ypjF* mutants. Colonies were inoculated onto semisolid agar plates and incubated at 30 °C for 20 hrs.

Next, we compared the proteomic rearrangements in each mutant. Two-dimensional plots revealed that the proteomic rearrangement by the ΔCP4-57 mutation correlated well with those of the Δ*ssrA* and Δ*ssrA*-*ypjF* mutants (**Fig. 2C**), but not with the Δ*intA*-*ypjF* mutant (**Fig. S2B**). These tendencies were also observed in *E. coli* cells in the stationary growth phase and those grown in a defined minimal medium (**Fig. S2C**). These results indicated that the proteomic rearrangement depends on the inactivation of the SsrA RNA, but not the loss of prophage-related genes in the CP4-57 region. The expression of SsrAΔU357 in the Δ*ssrA* mutant had no effect, excluding the possibility that the SsrAΔU357 molecule gains a novel biological function to rearrange the proteome (**Fig. S2D**).

We also examined whether the phenotypic alteration by the CP4-57 excision also depends on the switching of ribosome rescue pathways. Wang and colleagues reported that the CP4-57 excision affects several phenotypes, such as motility (28). Therefore, we introduced the mutations shown in **Fig. 2A** into the highly motile *E. coli* MG1655 strain, because the BW25113 strain is motility-impaired (36) (**Fig. S2E**). The proteomic rearrangements in MG1655 and its derivatives generated almost identical results to those of the BW25113 strains (**Fig. S2F**). All of the ΔCP4-57, Δ*ssrA* and Δ*ssrA*-*ypjF* mutants that lacked the *trans*-translation activity swam poorly (**Fig. 2D**), consistent with the previous study by Komine and colleagues, who reported the weakened motility phenotype of the Δ*ssrA* mutant (12), supporting our assumption. From these results, we concluded that the CP4-57 excision rearranges the proteome landscape and changes the motility phenotype by switching the ribosome rescue pathways.

### The absence of ribosome rescue-associated proteolysis of nonstop polypeptides triggers a dynamic rearrangement of the proteome

Both *trans*-translation and alternative rescue by ArfA commonly rescue stalled ribosomes, but only the former pathway induces the degradation of nonstop polypeptides. Accordingly, we assumed that the major determinant of the proteomic rearrangement by the CP4-57 excision would be the presence or absence of the degradation mechanism for the nonstop polypeptides between the two rescue pathways. To assess this, we again performed the SWATH-MS analysis for *E. coli* expressing the SsrA^DD^ or SsrA^His^ variant, which rescues the stalled ribosome but poorly induces the proteolysis of nonstop polypeptides (4, 8, 29, 37) (**Figs. 1B and 3A**). According to our hypothesis, we expected that the proteomic rearrangement in the SsrA^DD^ or SsrA^His^ strain would share some similarity with that in the Δ*ssrA* mutant.

*E. coli* expressing the SsrA^DD^ or SsrA^His^ mutant showed a certain degree of proteomic rearrangement, similar to that of the Δ*ssrA* mutant, relative to the wild-type SsrA-expressing strain (**Fig. S3A**). Furthermore, the proteomic rearrangements in these SsrA mutants correlated well with that of the Δ*ssrA* mutant (**Fig. 3B**). Actually, a moderate heat shock-like response was commonly induced in these proteolysis-deficient SsrA-expressing cells and the Δ*ssrA* cells (**Fig. S3B**). From these results, we concluded that the presence or absence of ribosome rescue-associated proteolysis activity is the major determinant of the proteomic rearrangement by the CP4-57 excision. The weakened motility phenotype of *E. coli* lacking the rescue-dependent proteolysis also supports this notion (38) (**Fig. 3C**).

**Fig. 3.**
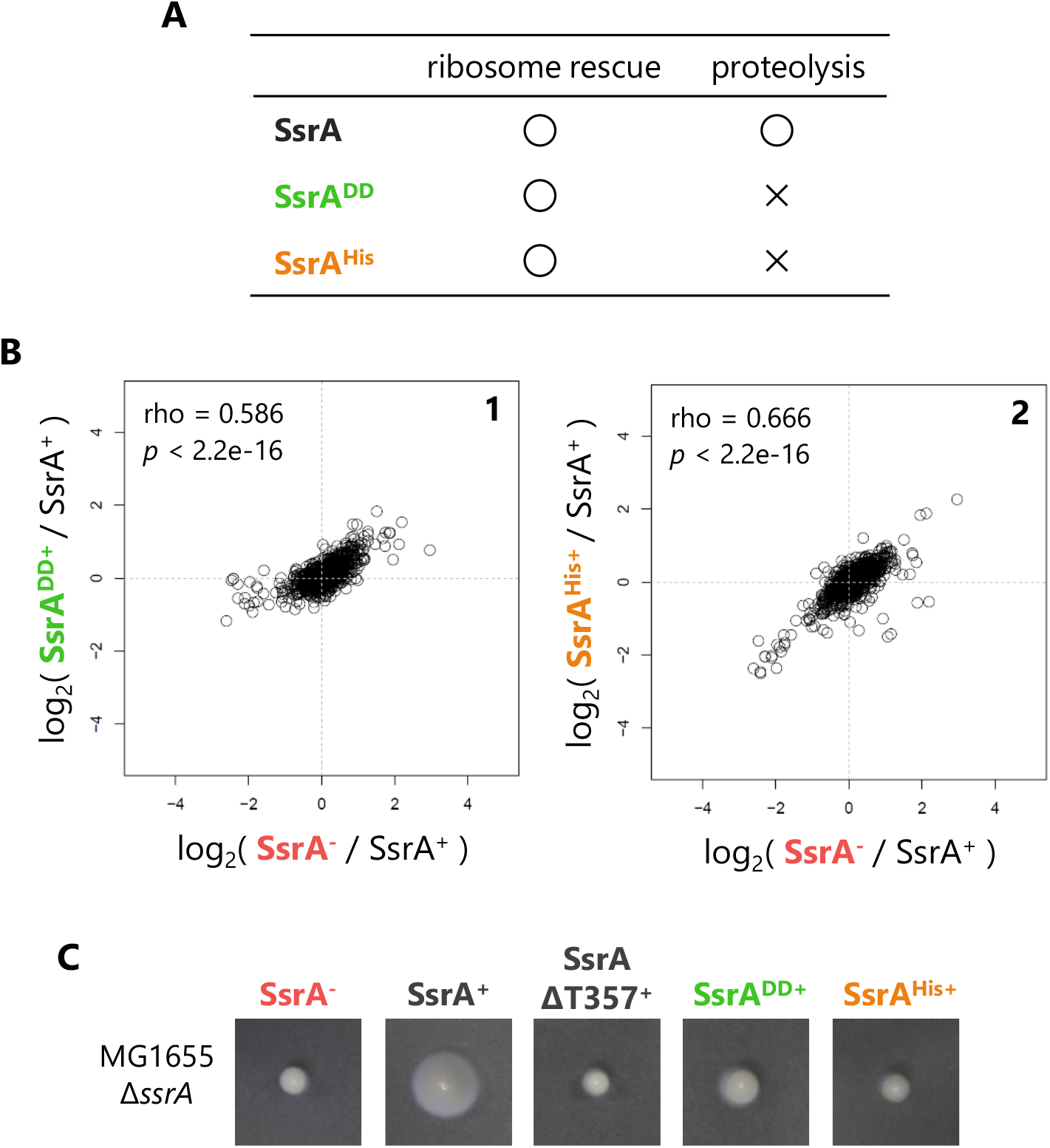
The absence of ribosome rescue-associated proteolysis of a nonstop polypeptide triggers the rearrangement of the cellular proteome. (A) Functional comparison of wild-type SsrA RNA and degradation tag mutants (SsrA^DD^ and SsrA^His^) that rescue the stalled ribosome but poorly induce the proteolysis of the released polypeptide (4, 8, 29, 37). The MG1655 Δ*ssrA* strain harboring pMW118 (vector control) and its derivatives expressing the SsrA variant were grown and analyzed as shown in **Fig. 2**. (B) Two-dimensional plots of the fold change values in each *E. coli* strain expressing SsrA or proteolysis-deficient mutants. Each plot is represented with Spearman’s rho and *P*-values calculated by Spearman’s rank correlation tests. (C) Swimming motility of the MG1655Δ*ssrA* strains harboring pMW118 and its derivatives, p-*ssrA*, p-*ssrA*ΔT357, p-*ssrA*^DD^ and p-s*srA*^His^. Colonies were inoculated onto semisolid agar plates and incubated at 30 °C for 20 hrs.

### ZntR expression is repressed in a *trans*-translation-dependent manner, but is independent of the degradation tag-induced proteolysis

In the course of the MS analyses, we noticed that the deficiency of the SsrA RNA reproducibly increased the expression of ZntR, a transcriptional regulator for the ZntA transporter that exports divalent metal ions such as Zn^2+^ (**Fig. 4A**). Interestingly, *zntR* is located just upstream of *arfA*, which is expressed as a nonstop mRNA (**Fig. 4B**). Due to the absence of a typical rho-independent terminator in the intergenic region between *zntR* and *arfA* (**Fig. S4A**), these genes are likely to be transcribed as a polycistronic mRNA. Accordingly, we hypothesized that the SsrA RNA-dependent repression of *zntR* is somehow coupled with the translation of the *arfA* nonstop mRNA. To assess this, the expression levels of ZntR with or without the following *arfA* ORF were examined by immunoblotting, using an N-terminal HA-tagged ZntR (**Fig. 4C**). When *zntR* was expressed as a monocistronic mRNA, the SsrA RNA had no impact on the expression of the ZntR protein (**Fig. 4D lanes 1-2**), confirming that ZntR is not a direct substrate of *trans*-translation. Disruption of the *arfA* promoter within the *zntR* ORF had no effect on the regulation of ZntR, excluding the possibility of premature transcription termination as in the case of *lacI* (39) (**Fig. S4B**). In the presence of SsrA RNA, the ZntR protein was poorly expressed from the *zntR-arfA* polycistronic mRNA (**Fig. 4D lanes 3-4**). In addition, a disruption of the initiation codon for the *arfA* ORF abolished the SsrA-dependent repression of ZntR (**Fig. 4D lanes 5-6**). From these results, we concluded that the translation of the nonstop *arfA* ORF induces the repression of ZntR in a *trans*-translation-dependent manner.

**Fig. 4.**
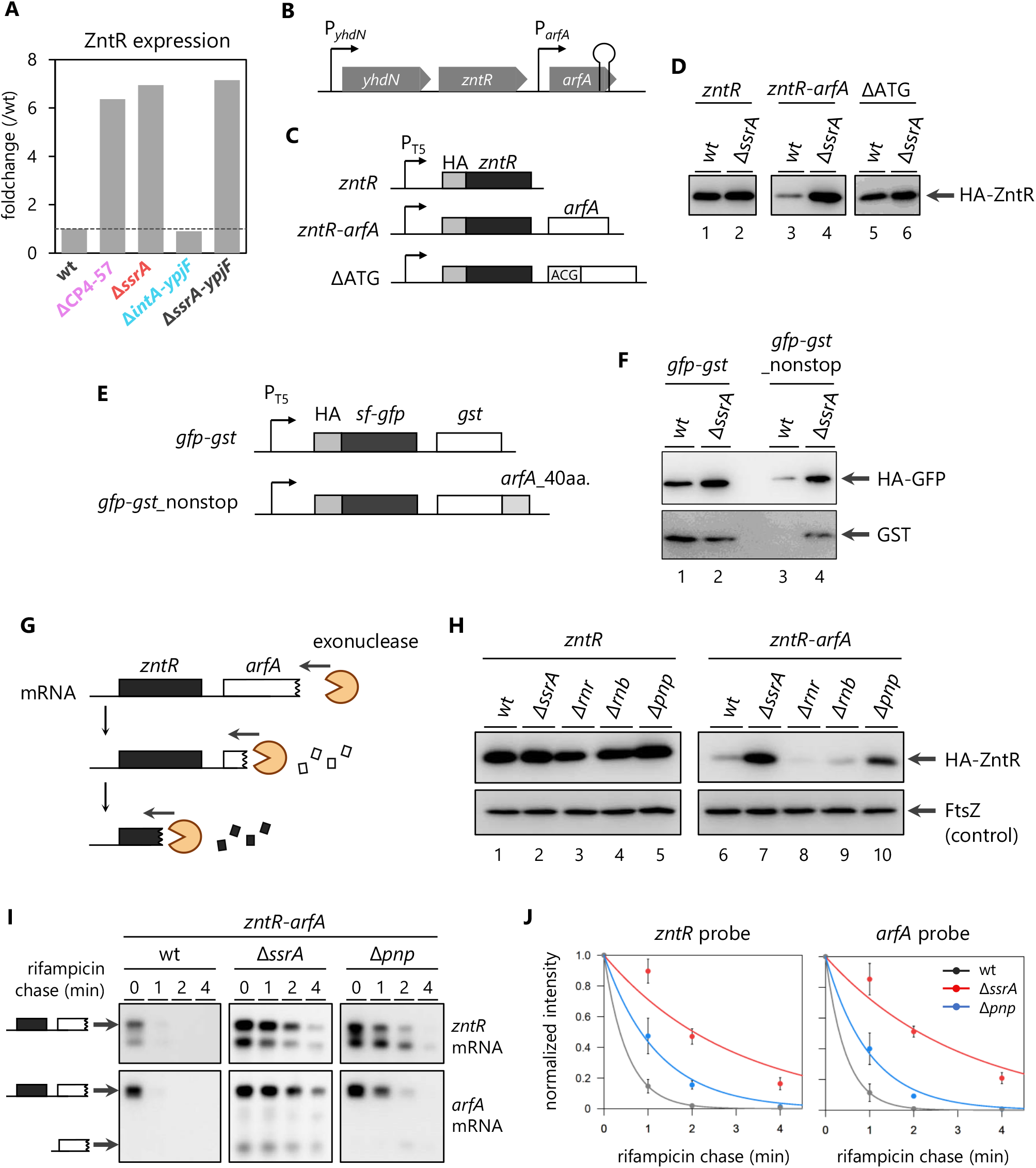
*zntR* mRNA is involved in the *trans*-translation-dependent degradation of *arfA* nonstop mRNA by the PNPase exonuclease. (A) Expression level of ZntR quantified by the SWATH-MS analyses in **Fig. 2**. The dashed line indicates a fold change value of 1. (B) Genetic structure of the *yhdN*-*zntR-arfA* locus. The transcription promoters of *yhdN* (P_*yhdN*_) and *arfA* mRNA (P_*arfA*_) and the intrinsic transcriptional terminator within the *arfA* ORF are schematically indicated. (C) Schematic representations of the N-terminally HA-tagged *zntR* and *zntR*-*arfA*. The initiation codon of *arfA* AUG was changed to ACG to deplete the translation of nonstop *arfA* mRNA (ΔATG). (D) Expression of HA-ZntR in the absence of SsrA RNA. The BW25113 (wt) and Δ*ssrA* strains harboring pOH012 (*zntR*, lanes 1 and 2), pOH008 (*zntR*-*arfA*, lanes 3 and 4) or pOH019 (ΔATG, lanes 5 and 6) were grown in LB medium until the OD_660_ reached 0.2-0.3. IPTG (100 µM) was then added to induce the expression of *zntR*, and cells were grown until the OD_660_ doubled. The cellular extracts were analyzed by western blotting using an anti HA-tag antibody. (E) Schematic representation of sfGFP-GST constructs. The entire sequences of the *zntR* and *arfA* ORFs are replaced by sfGFP (Superfolder GFP) and GST (glutathione-S transferase), respectively. The region encoding the C-terminal 40 amino acids of *arfA* was fused with the *gst* ORF to post-transcriptionally remove its stop codon (*gst*_nonstop). (F) Expression of sfGFP and GST in BW25113 (wt) or the Δ*ssrA* strain harboring pOH074 (*gfp*-*gst*, lanes 1 and 2) or pOH075 (*gfp*-*gst*_nonstop, lanes 3 and 4). Cellular extracts were prepared as shown in **Fig. 4D**, and analyzed by western blotting using an anti-GFP tag antibody (upper panel) and an anti-GST tag antibody (lower panel). (G) A working hypothesis for SsrA RNA-dependent ZntR repression: the *zntR* mRNA is involved in the exonucleolytic degradation of the *arfA* nonstop mRNA. (H) Expression of HA-ZntR in the mutants lacking one of the major exonucleases. BW25113 (wt) and its derivatives, Δ*ssrA*, Δ*rnr*, Δ*rnb* and Δ*pnp* strains harboring pOH090 (*zntR*, lanes 1-5) or pOH089 (*zntR*-*arfA*, lanes 6-10) were analyzed as in **Fig. 4D**. (I) Stability of the *zntR*-*arfA* polycistronic mRNA. Schematic labels indicate the *zntR*-*arfA* mRNA and the *arfA* mRNA. Wild-type, Δ*ssrA* andΔ*pnp* strains harboring pOH089 (*zntR*-*arfA*) were grown in LB medium until mid-log phase. The expression of the *zntR*-*arfA* mRNA was then induced by 1 mM IPTG for 15 min. At this point, 200 µg/ml of rifampicin was added to the culture. The total RNA was extracted from the cells at 0, 1, 2 and 4 min after rifampicin treatment. Total RNAs were analyzed by northern blotting using an anti-*zntR* mRNA probe (upper panel) and an anti-*arfA* mRNA probe (lower panel). (J) The quantified intensity of the *zntR*-*arfA* mRNA in (I). Error bars represent the SD values of three biological replicates.

The results described above indicated that the repression of ZntR is independent of the SsrA-driven proteolysis. To verify this further, we replaced the *zntR* and *arfA* ORFs with sfGFP (Superfolder GFP) and GST (glutathione S-transferase), respectively (**Fig. 4E**). The region encoding the C-terminal 40 amino acids of *arfA* was fused with the *gst* ORF to post-transcriptionally remove its stop codon (*gst*_nonstop). Translation of the *gfp*-*gst* polycistronic mRNA produced the sfGFP and GST proteins in a constant ratio, irrespective of the SsrA RNA (**Fig. 4F lanes 1-2**). In contrast, the expression of the sfGFP protein from the *gfp-gst*_nonstop mRNA, as well as that of the downstream GST, was significantly repressed in the presence of the SsrA RNA (**Fig. 4F lanes 3-4**). This result is quite consistent with the case of the *zntR*-*arfA* nonstop mRNA, confirming that the repression of ZntR is independent of its amino acid sequence and dependent on the context of the ORF followed by the nonstop ORF.

### *zntR* mRNA is involved in *trans*-translation-dependent degradation of *arfA* nonstop mRNA by exonuclease PNPase

*Trans*-translation stimulates the exonuclease degradation of nonstop mRNA (10). Therefore, we hypothesized that ZntR expression is repressed at the mRNA level; namely, the *zntR* mRNA is involved in the degradation of the *arfA* nonstop mRNA by the exonuclease(s) (**Fig. 4G**). Actually, the insertion of an artificial stem-loop into the intergenic region between *zntR* and *arfA* abolished the SsrA-dependent repression of ZntR, supporting this assumption (**Fig. S4C**). We evaluated the expression of ZntR in *E. coli* strains lacking one of the three major 3’→5’ exonucleases in *E. coli*. The monocistronic *zntR* mRNA was translated into the ZntR protein at almost identical levels in the wild-type and the Δ*ssrA* and exonuclease-lacking mutants (**Fig. 4H lanes 1-5**). In contrast, the *zntR*-*arfA* polycistronic mRNA allowed the synthesis of ZntR only in the Δ*ssrA* and Δ*pnp* mutants (**Fig. 4H lanes 7, 10**). This result indicated that PNPase, encoded by the *pnp* ORF, degrades the *zntR*-*arfA* mRNA in a *trans*-translation-dependent manner. PNPase is unable to degrade stable stem-loop structures in mRNA (40), consistent with the influence of the stem-loop insertion (**Fig. S4C**).

To obtain direct evidence, we next evaluated the stability of the *zntR*-*arfA* mRNA in the *E. coli* Δ*pnp* mutant. We stopped the synthesis of nascent mRNAs by the addition of rifampicin and extracted cellular RNA at the indicated time points. The mRNA molecules containing the *zntR* or *arfA* sequence were then individually probed by northern blotting. The monocistronic *zntR* mRNA was equivalently stable in all *E. coli* strains tested, again confirming that the *zntR* mRNA itself is not affected by *trans*-translation or PNPase (**Fig. S4D and S4E**). In contrast, the *zntR*-*arfA* polycistronic mRNA was extremely unstable in the wild-type strain, but significantly stabilized in the Δ*ssrA* and Δ*pnp* mutants (**Fig. 4I and 4J**). These results indicated that *trans*-translation triggers the PNPase-dependent degradation of the *zntR*-*arfA* mRNA so that the expression of ZntR is repressed in the presence of the SsrA RNA. Consistent with this notion, we reanalyzed the proteomic datasets of the SsrA variants (**Fig. 3**) and found that the SsrA^DD^ mutant also repressed the ZntR expression (**Fig. S4F**). The SsrA^DD^ variant is defective in the proteolysis of nonstop polypeptides, but efficiently promotes the degradation of nonstop mRNA as well as the wild-type SsrA RNA (9). From these results, we concluded that *trans*-translation regulates the ZntR expression in a proteolysis-independent and exonucleolytic degradation-dependent manner.

## Discussion

Previous studies of ArfA, the SsrA RNA-dependent tight repression and the cooperative action with RF2 to release stalled ribosomes have been performed from a variety of perspectives, including biochemical and structural analyses (20, 21, 41–44). However, the situations in which the ArfA/RF2 pathway is activated to compensate for the ribosome rescue function, and the impact on the cellular proteome when the ribosomal rescue pathway is actually switched to the alternative rescue pathway, have not been investigated.

In this study, we demonstrated that the excision of the CP4-57 prophage, located downstream of the *E. coli ssrA* gene, functions as a phage regulatory switch to inactivate the SsrA RNA and to simultaneously activate the ArfA/RF2 ribosome rescue system (**Fig. 1**). The excision of CP4-57 deletes the U357 residue of SsrA, which forms a G·U wobble base pair in the acceptor stem of tmRNA (28). Biochemical and structural analyses of the wobble base pair have shown its importance for the recognition of alanyl-tRNA synthetase (45–48), indicating that SsrAΔU357 has an aminoacylation defect. This inactivation of the SsrA RNA rearranged the intracellular proteome of *E. coli* and also altered the phenotype (**Fig. 2**). Furthermore, the similarity between the proteomic rearrangements among Δ*ssrA* and the proteolysis-deficient SsrA mutants indicates that the accumulation of the nonstop polypeptide is the major determinant for the reorganization of the *E. coli* proteome (**Fig. 3, summarized in Fig. 5A**).

**Fig. 5.**
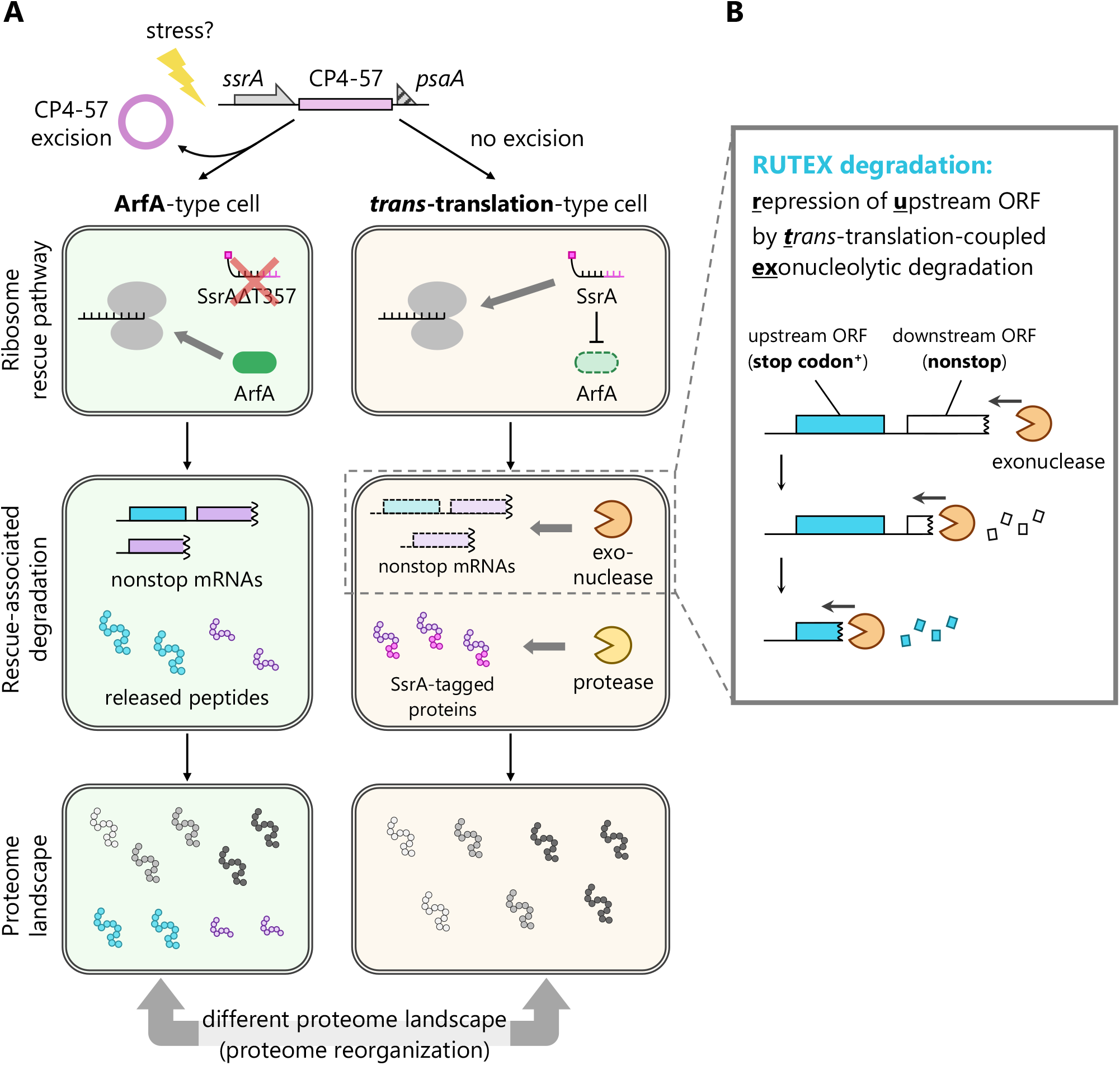
Proposed mechanism of the *E. coli* proteome rearrangement by the CP4-57 excision. (A) Excision of the CP4-57 prophage from the *E. coli* genome is triggered by various environmental stresses. This DNA rearrangement also induces a single nucleotide deletion in *E. coli ssrA*, which impairs the *trans*-translation activity and subsequently activates the ArfA/RF2 ribosome rescue pathway back-to-back. This switching results in the accumulation of nonstop polypeptides that are usually repressed by *trans*-translation-coupled proteolytic and nucleolytic degradation, thus reorganizing the proteome landscape. (B) Schematic of the repression of upstream ORF by ***t****rans*-translation-coupled exonucleolytic (RUTEX) degradation.

In addition to *S. oneidensis, E. coli* and other γ-proteobacteria possess phage regulatory switches that irreversibly shut off the function of the SsrA RNA (**Table 1 and Fig. S1B**). Since all bacterial species with the prophage-dependent SsrA-inactivation mechanism also have ArfA as an alternative rescue mechanism, the prophage excision would switch the ribosome rescue pathway, as in *E. coli*. However, the excision of other *ssrA*-adjacent prophages does not always modify the sequence of the *ssrA* gene, even though their host cells possess *arfA* (*e*.*g*., *Salmonella Typhimurium* DT204 and *E. coli* CFT073, **Table 1** (31, 49)). Therefore, the phage regulatory switches that regulate the ribosome rescue functions among different bacterial species were probably acquired independently after the phage integration.

At first glance, the inactivation of *trans*-translation is merely a defective event for the survival of bacterial cells, as reported in previous studies (12, 50–52). However, the excision of CP4So in *S. oneidensis* or CP4-57 in *E. coli* is induced by various environmental stresses and contributes to survival in certain situations (22, 28). In this context, it is noteworthy that the excisions of CP4So and CP4-57 contributed to biofilm formation through a stochastic differentiation of the bacterial subpopulations. Taken together with the previous report that biofilm formation requires a transition from a motile to a non-motile state (53), the non-motile ΔCP4-57 cells would form the early stage of the biofilm as the subpopulation appeared from the motile wild-type cells. In general, the excision of prophages including CP4-57 in *E. coli* is induced stochastically in a small part of the bacterial population under certain stress conditions, consistent with this assumption (27). However, the determinants for the frequency of the prophage excision and the overall landscape of the environmental stresses that trigger the excision are not fully understood. Further analyses will be required to elucidate the biological significance of *trans*-translation inactivation by CP4-57 excision.

Using proteomics analyses as a starting point, we revealed that the expression of ZntR is repressed by PNPase-dependent degradation, which is independent of the sequence features of *zntR* but dependent on the following nonstop *arfA* ORF (**Fig. 4**). To our knowledge, previous related studies mainly focused on the translation of the stop codon-less ORF itself, and thus the influence of *trans*-translation on the neighboring genes such as *zntR* would have been overlooked. Accordingly, our findings on *zntR* expand the repertoire of *trans*-translation-dependent gene regulation and differ from the canonical substrates of SsrA RNA in the following points: (i) *zntR* (and maybe other genes) is regulated through *trans*-translation even though its mRNA maintains the stop codon, and (ii) the expression of ZntR is repressed by mRNA degradation, and is independent of the degradation tag-dependent proteolysis. Here we propose to define this <u>**r**</u>epression of <u>**u**</u>pstream ORF by ***<u>t</u>****rans*-translation-coupled <u>**ex**</u>onucleolytic degradation as RUTEX degradation (**Fig. 5B**).

From another viewpoint, RUTEX degradation determines the stability of the target mRNA by the translation status of the 3’ downstream region, irrespective of the translation of its own ORF. Expanding this notion, we assume that RUTEX degradation can also regulate an apparent monocistronic mRNA carrying an intact ORF. For example, translation of the 3’ UTR by stop codon readthrough or frameshifting may induce *trans*-translation at the regular 3’ end of the mRNA and the following mRNA degradation by exonuclease(s). Thus, it is possible that RUTEX degradation extends the role of *trans*-translation beyond the degradation of aberrant products. RUTEX degradation might define the limit of translation frequency per each mRNA in a cooperative action with the stochastic occurrence of noncanonical translation, such as stop codon readthrough and frameshifting at the C-terminus of the ORF. Our assumption contrasts with the well-studied concept that the translation of upstream ORFs (uORFs, or leader peptides) determines the translation frequency of the downstream ORF or mRNA expression (54). Interestingly, several groups reported that the translation of the 3’ UTR also regulates the translation frequency of the main ORF in eukaryotes (55, 56). RUTEX degradation might play a similar role in prokaryotes.

We found that the *zntR*-*arfA* mRNA is primarily degraded by PNPase (**Fig. 4H-J**). Herzel and colleagues recently reported that the amount of the *arfA* transcript increases in a Δ*pnp* strain (57), in agreement with our results. However, the contribution of PNPase disagrees with a previous finding that RNase R is responsible for the degradation of nonstop mRNA (10). We expect that one of the determinants for this divergence is the presence or absence of the secondary structure at the 3’ end of the mRNA. In detail, the expression of the artificial lambda cI nonstop mRNA is achieved by the insertion of the *trpA* rho-independent terminator within the cI ORF (10), whereas the deletion of the *arfA* stop codon is dependent on the endonucleolytic cleavage by RNase III, thus removing the stem-loop structure (20, 21). PNPase is unable to degrade mRNAs that form a stable stem-loop (40), while in contrast, RNase R exerts its exonuclease activity regardless of the stem-loop (58, 59). According to these previous findings and our results, the degradation of nonstop mRNAs is not likely to be uniform, as previously considered. Rather, we assume that exonucleases such as RNase R, PNPase and RNase II cooperatively degrade a large variety of nonstop mRNAs, and the preference for each substrate depends on the shape of the 3’ end of each nonstop mRNA. The *zntR*-*arfA* mRNA was relatively stable in the Δ*pnp* strain but more stable in the Δ*ssrA* strain, probably reflecting the contribution of other exonuclease(s) in the absence of PNPase (**Fig. 4J**). The existence of multiple exonucleases in bacterial species would contribute to the robust degradation of the aberrant mRNAs. However, the means by which *trans*-translation promotes PNPase-dependent degradation and the criteria of the exonuclease selection for the degradation of each mRNA are still unknown. Further analyses will be required to elucidate the detailed mechanism of the exonuclease recruitment and clarify the overall picture of aberrant mRNA quality control in bacterial species.

## Author Contributions

H.O., T.N., and Y.C. performed experiments; H.O., T.N., H.T., and Y.C. conceived the study, designed experiments and analyzed the results; H.T., and Y.C. supervised the entire project; H.O., H.T., and Y.C. wrote the manuscript.

## Acknowledgments

We thank Eri Uemura for technical support and the Bio-support Center at Tokyo Tech for DNA sequencing. This work was supported by MEXT Grants-in-Aid for Scientific Research (Grant Numbers JP26116002, JP18H03984, JP20H05925 to HT, 17K15062, 19K16038 to YC) and a grant from the Ohsumi Frontier Science Foundation to YC.

## Data sharing plan

The MS-datasets obtained in this study will be deposited in jPOST repository (https://repository.jpostdb.org).

Sequence files of the plasmids constructed in this study will be deposited in Mendeley Data repository (https://data.mendeley.com).

## Materials and Methods

### *E. coli* strains, plasmids, and primers

*E. coli* strains, plasmids and primers used in this study are listed in **Tables S1, S2 and S3**, respectively. Phage P1-mediated transduction was used to introduce the *arfA, rnr, rnb* and *pnp* mutations from the Keio collection JW3253, JW5741, JW1279 and JW5851, respectively (60). The Δ*ssrA*, Δ*intA*-*ypjF* and Δ*ssrA*-*ypjF* mutations were introduced by homologous recombination (61) with a chloramphenicol resistance cassette flanked by FRT (FLP recognition target) sites. The DNA fragments for introducing the Δ*ssrA*, Δ*intA*-*ypjF* and Δ*ssrA*-*ypjF* mutations were amplified using the primer pairs ON001 and ON002, ON027 and ON028, and ON001 and ON028, respectively, with pCGT1 as the template. Each purified DNA was electroporated into *E. coli* strain BW25113 or MG1655 harboring pKD46, and the transformants resistant to 20 µg/ml chloramphenicol were stored as the Δ*ssrA*, Δ*intA*-*ypjF* and Δ*ssrA*-*ypjF* mutants, respectively.

BW25113 or MG1655 carrying pCY2794 (derivative of pKD46 with *alpA* (26) under the control of the P_BAD_ promoter) was grown in LB medium containing 0.2 % arabinose and 100 µg/ml ampicillin at 30 °C for 2 hrs to induce the excision of CP4-57 from the *E. coli* chromosome. A 2 µl portion of the culture was spread onto an LB agar plate and incubated at 37 °C overnight to remove pCY2794. Isolated colonies were streaked onto LB agar plates with or without 100 µg/ml ampicillin to monitor the loss of pCY2794. Complete removal of CP4-57 was confirmed by PCR, using ON003 and ON225. Plasmids were constructed using standard cloning procedures and Gibson assembly (62). Detailed schemes for plasmid construction are summarized in **Table S2**.

### Bacterial growth and motility assay

*E. coli* cells were grown in LB medium or MOPS minimal medium (63) at 37 °C unless otherwise noted. LB containing 1.5 % (w/v) agar was used to isolate *E. coli* colonies. Appropriate antibiotics were added to the media. Bacterial growth in liquid medium was monitored by measuring the OD_660_. Cell motility was examined with semisolid agar plates (0.5 % peptone, 0.3 % yeast extract, 0.3 % agar). Colonies were inoculated onto the plates and incubated at 30 °C for 20 hrs.

### Western blotting analysis

Cell cultures were withdrawn and mixed with an equal volume of 10% of TCA. After standing on ice for at least 10 min, the samples were centrifuged at 15,000 rpm for 3 min at 4 °C, and the supernatant was removed by aspiration. Precipitates were washed with acetone by vigorous mixing, centrifuged again, and dissolved in 1× SDS sample buffer (62.5 mM Tris-HCl, pH 6.8, 5 % 2-mercaptoethanol, 2 % SDS, 5 % sucrose, and 0.005 % bromophenol blue) by vortexing for 15 min at 37 °C. Prepared cellular extracts were separated by 13 or 15 % SDS-PAGE and subsequently transferred to an Immobilon-P^SQ^ Membrane (Millipore). The membrane was blocked by 1 % (w/v) skim milk in TBS-Tween (20 mM Tris-HCl, pH 7.5, 150 mM NaCl, 0.1 % Tween-20) at room temperature for 1 hr. The membrane was then incubated with TBS-Tween containing 1 % (w/v) skim milk and an antibody (1/10,000 dilution) at room temperature for 1 hr. Plasmid-borne ArfA, ZntR, sfGFP, GST and endogenous FtsZ were detected by an Anti 6×Histidine Monoclonal Antibody (9C11, Wako), Monoclonal Anti-HA clone HA-7 (Sigma-Aldrich), Anti-Green Fluorescent Protein Monoclonal Antibody (mFx75, Wako), GST·Tag Monoclonal Antibody (Sigma-Aldrich) and Anti-FtsZ antibody (a gift from Dr. Shinya Sugimoto at Jikei Medical University), respectively. HRP-conjugated anti-mouse IgG (Sigma-Aldrich) and HRP-conjugated anti-rabbit IgG (Sigma-Aldrich) were used as secondary antibodies. Images were visualized and analyzed by a LAS4000 LuminoImager (Fujifilm).

### Northern blotting analysis

*E. coli* cells were grown in LB medium until the OD_660_ reached ∼0.5. The expression of *zntR* mRNA or *zntR*-*arfA* mRNA was then induced by 1 mM IPTG for 15 min. At this point, 200 µg/ml of rifampicin was added to the culture. The culture was harvested at 0, 1, 2 and 4 min after the addition of rifampicin. Total RNA was prepared using the Tripure Isolation Reagent (Roche), according to the supplier’s instructions. RNA samples were separated by 1.5 % denaturing agarose electrophoresis, transferred onto BrightStar-Plus Positively Charged Nylon Membranes (Invitrogen), and hybridized with biotinylated oligonucleotides (Integrated DNA Technologies) complementary to the *zntR* or *arfA* mRNA, as shown in **Table S4**. Hybridization experiments were performed using a NorthernMax kit (Ambion) and a Chemiluminescent Nucleic Acid Detection Module (ThermoFisher Scientific) according to the suppliers’ instructions. Images were visualized and analyzed by a LAS4000 LuminoImager (Fujifilm).

### Sample preparation for proteomic analysis

Cells were grown in LB medium until the OD_660_ reached ∼0.5. After the cells were harvested, they were washed and resuspended with PBS buffer (137 mM NaCl, 8.1 mM Na_2_HPO_4_, 2.68 mM KCl, 1.47 mM KH_2_PO_4_, pH 7.4). The suspension was mixed with an equal volume of 10% of TCA. After standing on ice for at least 10 min, the samples were centrifuged at 15,000 rpm for 5 min at 4 °C, and the supernatant was removed by aspiration. Precipitates were washed twice with acetone, by vigorous mixing. Sample preparation for LC-MS/MS was basically performed according to the previous study (64) with some modifications. Proteins were dissolved in PTS solution (12 mM sodium deoxycholate, 12 mM sodium lauryl sulfate, 100 mM Tris-HCl, pH 9.0) and the concentration of the lysate was determined by using a Pierce BCA Protein Assay Kit (ThermoFisher Scientific). Subsequently, 50 µg of total protein at a concentration of 1 µg/µl was processed according to the following procedure. First, the lysate was reduced by a treatment with 20 mM dithiothreitol (DTT) at room temperature for 30 min, and then alkylated with 50 mM iodoacetamide in the dark at room temperature for 20 min. The protein mixture was then diluted 5-fold with 50 mM ammonium bicarbonate. For the limited digestion of denatured proteins into peptide fragments, 0.5 µg of Trypsin/Lys-C Mix (Promega, U.S.A.) was added, and the mixture was incubated at room temperature for 3 hrs. Subsequently, 1.0 µg of Trypsin/Lys-C Mix was again added and incubated at 37 °C overnight. After the digestion, an equal volume of ethyl acetate and 0.5 % trifluoroacetic acid (final concentration) was added. The mixture was shaken vigorously for 2 min and then centrifuged at 15,700 ×g for 2 min. The upper ethyl acetate layer was discarded, and the solvent was removed by a centrifugal evaporator. The residual pellet was redissolved in 600 µl of 0.1 % TFA and 2 % acetonitrile, and desalted as follows. The solution was applied to a StageTip composed of an SDB-XC Empore disk (3M, U.S.A.), equilibrated with 0.1 % TFA and 2 % acetonitrile, washed with 0.1 % TFA and 2 % acetonitrile, and eluted with 0.1 % TFA and 80 % acetonitrile. The solvent was removed by a centrifugal evaporator, and the residual peptides were redissolved in 120 µl of 0.1 % TFA and 2 % acetonitrile. This solution was centrifuged at 20,000 ×g for 5 min, and a 100 µl portion of the supernatant was collected and subjected to the LC-MS/MS measurement.

### LC-MS/MS-based proteomic analysis

The LC-MS/MS measurements (33) (SWATH-MS acquisition) were conducted with an Eksigent NanoLC Ultra and TripleTOF 4600 tandem-mass spectrometer or an Eksigent nanoLC 415 and TripleTOF 6600 mass spectrometer (AB Sciex, U.S.A.). The trap column used for nanoLC was a 5.0 mm × 0.3 mm ODS column (L-column2, CERI, Japan) and the separation column was a 12.5 cm × 75 μm capillary column packed with 3 μm C18-silica particles (Nikkyo Technos, Japan). The detailed settings for the LC-MS/MS measurements are shown in **Table S5**. The SWATH acquisition was performed three times for each sample. Data analysis of the SWATH acquisition was performed using the DIA-NN software with default settings (65). The library for SWATH acquisition was obtained from the SWATH atlas (http://www.swathatlas.org, the original data are in Midha *et al*., 2020 (66)). Only the proteins detected in all three measurements for both samples were used for the fold change calculation. The obtained protein intensities were averaged by using an in-house R script. The *P*-value was determined by Welch’s t-test and corrected by the Benjamini-Hochberg method for multiple comparisons, using the “p.adjust” function in R (for Windows, version 4.1.2).

