## supplementary figures for "Novel *trans*-translation-associated gene regulation revealed by prophage excision-triggered switching of ribosome rescue pathway"

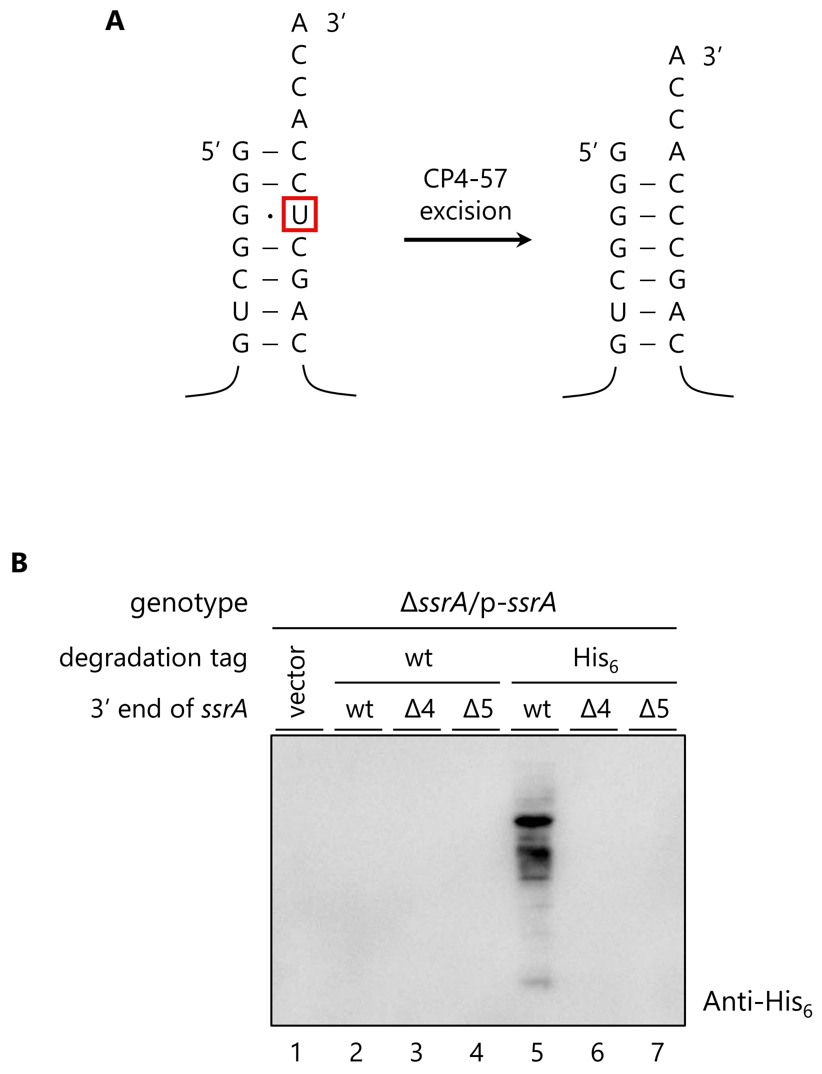


**Fig. S1.**

(A) The acceptor stem-loop structure at the 3' end of *E. coli* SsrA RNA and the SsrAΔU357 mutant. CP4-57 excision deletes the T357 residue of the *ssrA* gene, which forms a G·U wobble base pair in the acceptor stem in the tRNA-like domain.

(B) Inactivation of $SsrA^{His}$ by prophage excision-introduced mutations in other bacterial species. The *E. coli* Δ*ssrA* strain harboring pMW118 (vector control, lane 1) and its derivatives, p-*ssrA* (lane 2), p-*ssrA*Δ4 (induced by the excision of Ype11X in *Yersinia pestis* CO92, which deletes four nucleotides at the 3' end, lane 3), p-*ssrA*Δ5 (induced by the excision of Stm27X in *Salmonella enterica* serovar Typhimurium LT2, Eco48X in *Escherichia coli* RS218 and Oi108 in *Escherichia coli* O157:H7 EDL933, which deletes five nucleotides at the 3' end, lane 4), p-$ssrA^{His}$ (lane 5), p-$ssrA^{His}$Δ4 (lane 6) and p-$ssrA^{His}$Δ5 (lane 7) were grown in LB medium until mid-log phase. Cellular extracts were fractionated by SDS-PAGE and analyzed by western blotting using an anti-$His_6$ antibody.

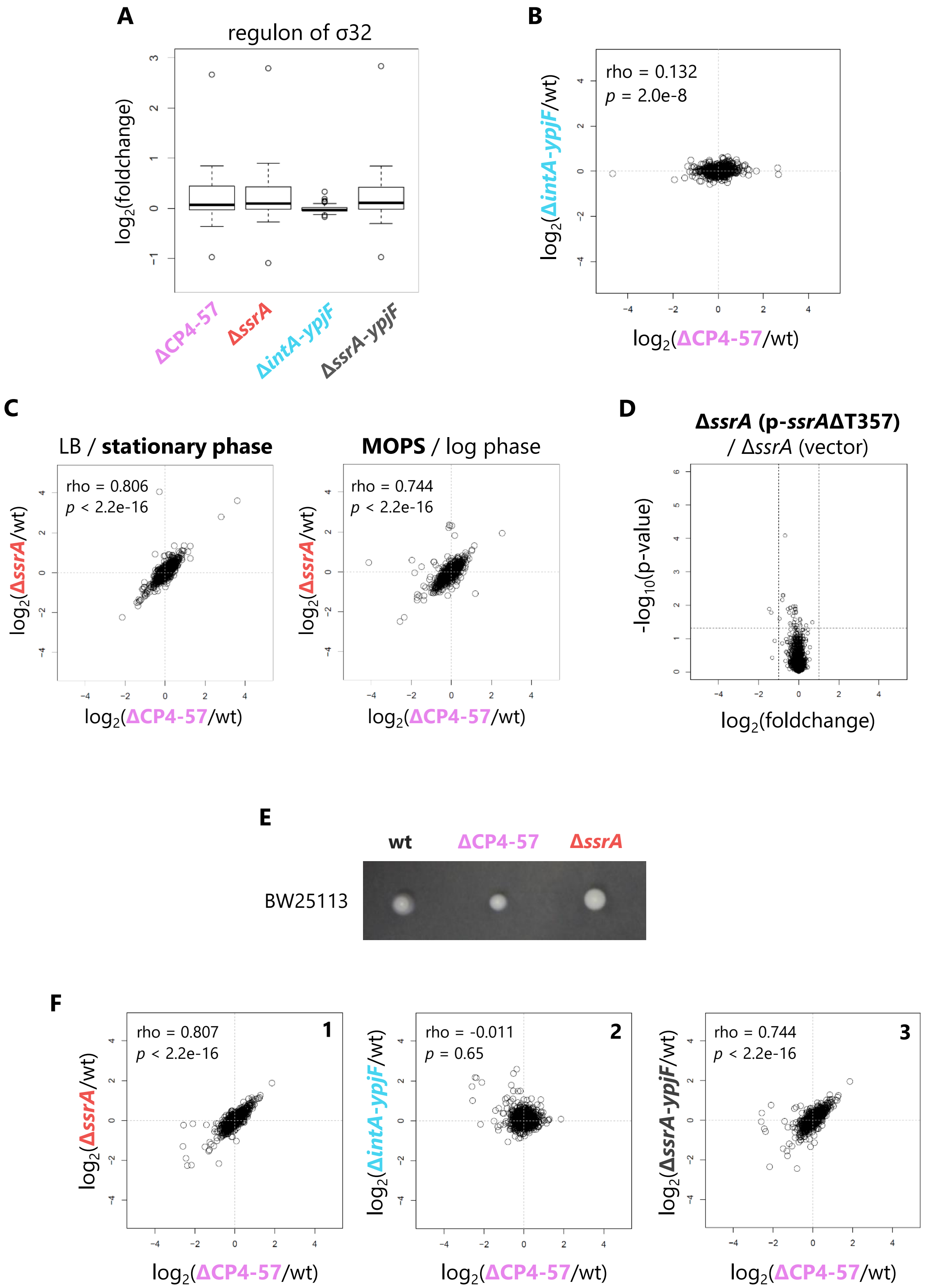


**Fig. S2.**

(A) Fold change values of the heat shock regulons (σ32) were extracted from the results in Fig. 2, and represented as a boxplot.

(B) Two-dimensional plots of the fold change values in the *E. coli* BW25113 ΔCP4-57 (horizontal axis) and Δ*intA-ypjF* strains (vertical axis). The plot is represented with Spearman's rho and *P*-values calculated by Spearman's rank correlation tests.
(C) Two-dimensional plots comparing the proteomic rearrangements in the BW25113 ΔCP4-57 and BW25113 Δ*ssrA* strains in LB medium at the stationary growth phase (left) or in MOPS minimal medium at the mid-log growth phase (right). Cells were grown at 37 °C in the indicated medium and then subjected to the SWATH-MS analysis, as shown in **Fig. 2**. Each plot is represented with Spearman's rho and *P*-values calculated by Spearman's rank correlation tests.
(D) A volcano plot showing the poor proteomic change in the Δ*ssrA* strain expressing SsrAΔU357, compared to the Δ*ssrA* strain harboring an empty vector. Fold change and *P*-values of each protein are represented by dots and plotted according to its $\log_2$ fold change on the horizontal axis and *P*-value on the vertical axis. The lines indicate a *P*-value of 0.05 and 2-fold change.
(E) Swimming motility of BW25113 wild-type strain and its derivatives, ΔCP4-57 and Δ*ssrA* mutant. Colonies were inoculated onto semisolid agar plates and incubated at 30 °C for 20 hrs.
(F) Two-dimensional plots of the fold change values in the *E. coli* MG1655 mutant indicated below and on the side of the graph. Each plot is represented with Spearman's rho and *P*-values calculated by Spearman's rank correlation tests.

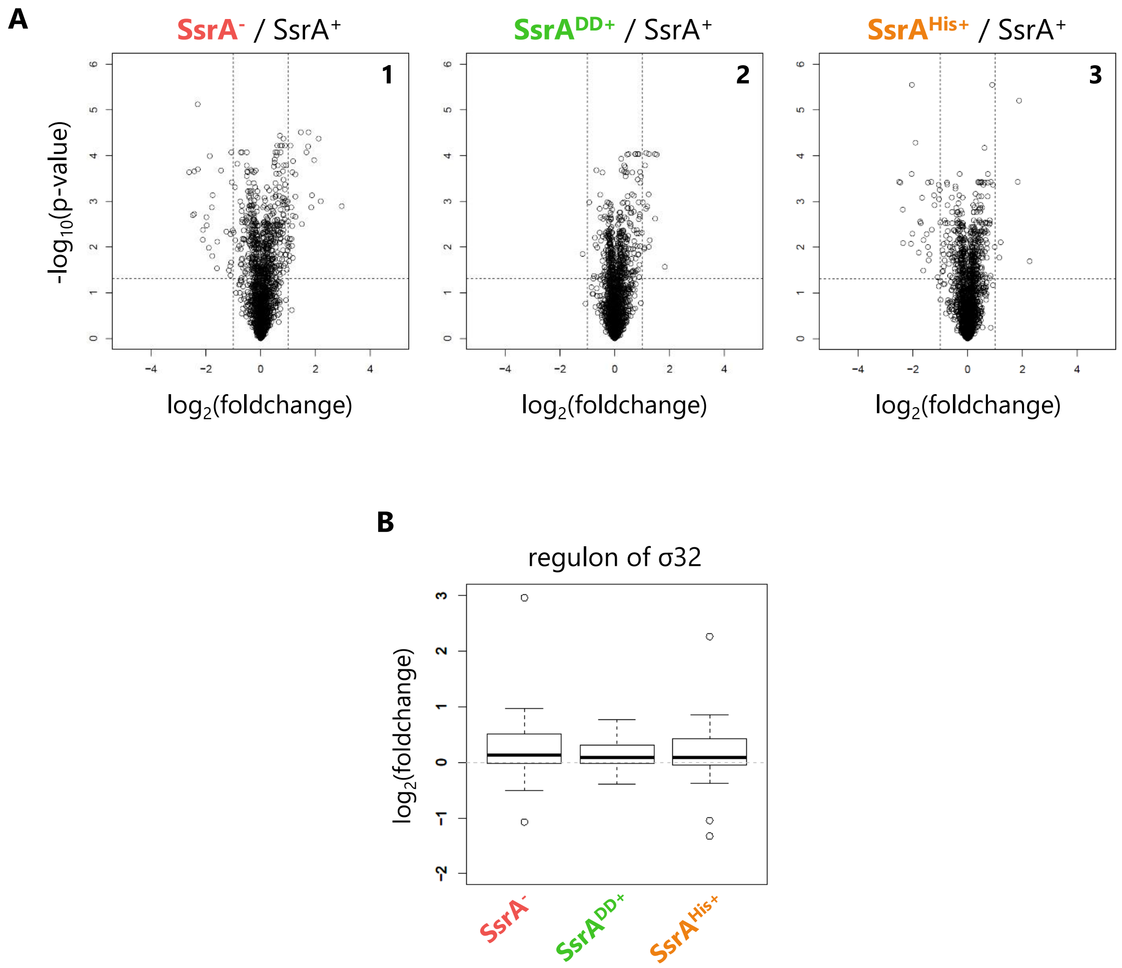


**Fig. S3.**

(A) Proteomic rearrangements in proteolysis-deficient SsrA-expressing strains. The fold change relative to the wild-type SsrA expressing strain and the *P*-value of each protein in the Δ*ssrA* strain harboring pMW118 (panel 1) and its derivatives, p-*ssrA*$^{DD}$ (panel 2) and p-*ssrA*$^{His}$ (panel 3), are represented by volcano plots, as shown in **Fig. 2**.

(B) Fold change values of the heat shock regulons (σ32) were extracted from the results in **Fig. 3**, and are represented by boxplots.

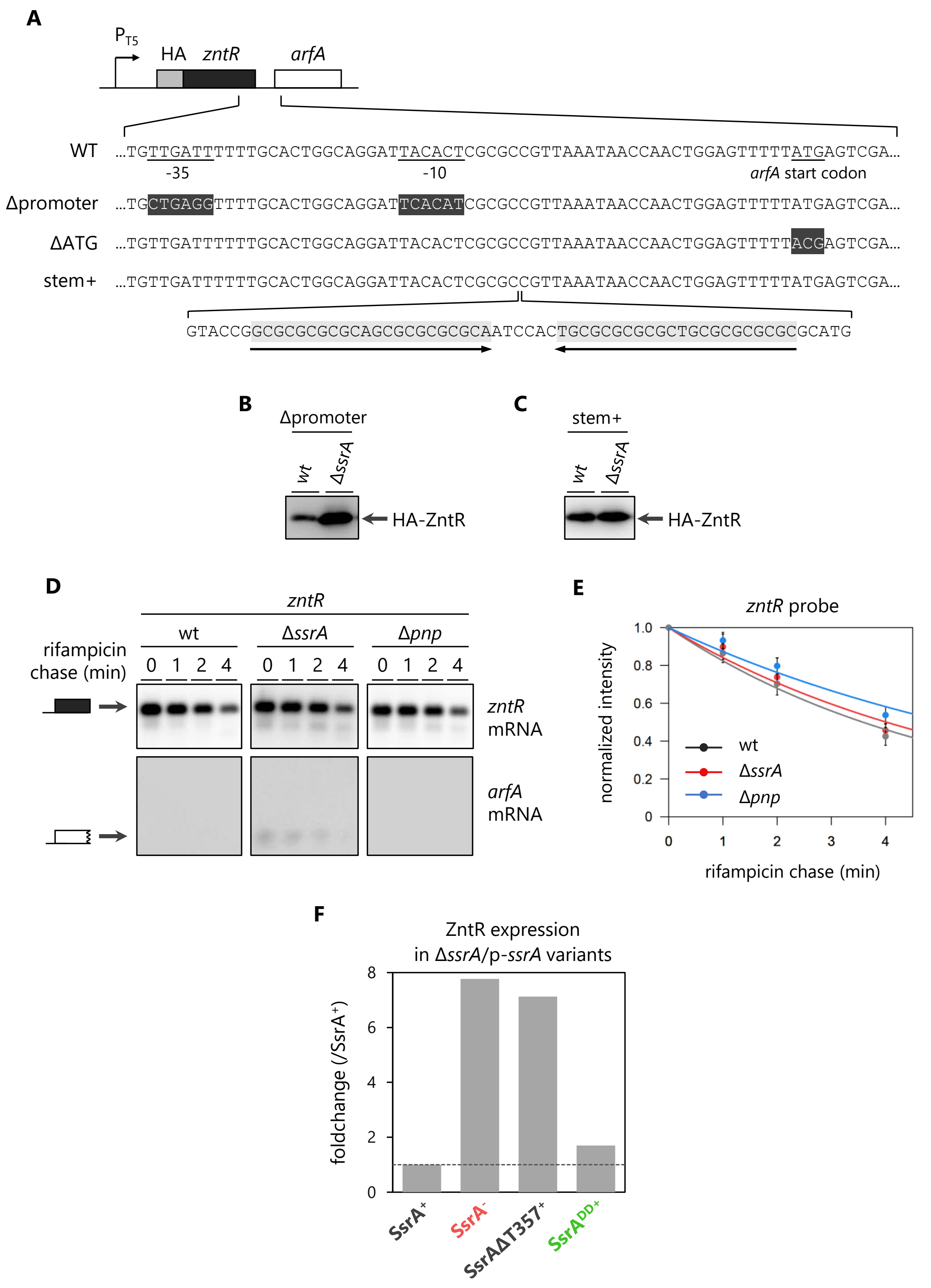


**Fig. S4.**

(A) Schematic representation of the mutations introduced into the *zntR*-*arfA* operon: WT (wild-type), ΔATG (disruption of the initiation codon of *arfA*), Δpromoter (disruption of *arfA* promoter) and stem+ (insertion of an artificial stem-loop between *zntR* and *arfA,* indicated by arrows).

(B) Expression of HA-tagged ZntR from *zntR*-*arfA* lacking the *arfA* promoter. BW25113 and the Δ*ssrA* strain harboring pOH020 (*zntR*-*arfA* Δpromoter) were grown in LB until the $OD_{660}$ reached 0.2-0.3. IPTG (100 μM) was then added to induce the expression of *zntR*, and cells were grown until the $OD_{660}$ doubled. Cellular extracts were prepared and analyzed as in **Fig. 4D**.
(C) Expression of HA-tagged ZntR from *zntR*-*arfA* with the insertion of an artificial stem-loop. BW25113 and the Δ*ssrA* strain harboring pOH021 (*zntR*-*arfA* stem+) were grown and analyzed as in **Fig. 4D**.
(D) Stability of the *zntR* monocistronic mRNA. Schematic labels indicate the *zntR* mRNA (filled box) and the nonstop *arfA* mRNA (open box). The wild-type, Δ*ssrA* strain andΔ*pnp* strain harboring pOH090 (*zntR*) were grown in LB medium until mid-log phase. The expression of *zntR* mRNA was then induced by 1 mM IPTG for 15 min. At this point, 200 μg/ml of rifampicin was added to the culture. The total RNA was extracted from the cells at 0, 1, 2 and 4 min after rifampicin treatment. Total RNAs were analyzed by northern blotting using an anti-*zntR* mRNA probe (upper panel) and an anti-*arfA* mRNA probe (lower panel).
(E) Quantified intensity of the *zntR* mRNA in **Fig. S4D**. Error bars represent the SD values of three biological replicates.
(F) Expression levels of ZntR in SsrA variant-expressing cells, extracted from the SWATH-MS analyses in **Fig. 3**. The dashed line indicates a fold change value of 1.
